# Excess of rare missense variants in hearing loss genes in sporadic Meniere disease

**DOI:** 10.1101/393322

**Authors:** Alvaro Gallego-Martinez, Teresa Requena, Pablo Roman-Naranjo, Jose A. Lopez-Escamez, Meniere Disease Consortium (MeDiC)

**Author notes:** Correspondence: Dr. Jose A. Lopez-Escamez, Otology & Neurotology Group CTS 495, Department of Genomic Medicine, GENYO. Centre for Genomics and Oncological Research: Pfizer/University of Granada/Andalusian Regional Government, PTS Granada, Avenida de la Ilustración, 114, 18016 Granada SPAIN.

## Abstract

Meniere’s disease (MD) is a clinical spectrum of rare disorders characterized by vertigo attacks, associated with sensorineural hearing loss (SNHL) and tinnitus involving low to medium frequencies. Although it shows familial aggregation with incomplete phenotypic forms and variable expressivity, most cases are considered sporadic. The aim of this study was to investigate the burden for rare variation in SNHL genes in patients with sporadic MD.

We conducted a targeted-sequencing study including SNHL and familial MD genes in 890 MD patients to compare the frequency of rare variants in cases using three independent public datasets as controls.

Patients with sporadic MD showed a significant enrichment of missense variants in SNHL genes that was not found in the controls. The list of genes includes *GJB2, USH1G, SLC26A4, ESRRB* and *CLDN14*. A rare synonymous variant with unknown significance was found in the *MARVELD2* gene in several unrelated patients with MD.

There is a burden of rare variation in certain SNHL genes in sporadic MD. Furthermore, the physical interaction of specific gene variants at protein level can explain the additive effect of rare variants in different genes in MD. This study will contribute to design a gene panel for the genetic diagnosis of MD.

## Introduction

Meniere’s disease (MD, MIM 156000) is a chronic disorder of the inner ear characterized by episodes of vertigo, associated with low to middle frequency sensorineural hearing loss (SNHL), tinnitus and aural fullness (Lopez-Escamez et al. 2015). The disorder produces an accumulation of endolymph in the membranous labyrinth, and it may affect both ears in 25-40% of patients (termed bilateral MD) and most of cases are considered sporadic (Paparella and Griebie 1984). However, heterogeneity in the phenotype is observed and some patients may have co-morbid conditions such as migraine or systemic autoimmune disorders (Gazquez et al. 2011; Caulley et al. 2017). This phenotypic spectrum can make the clinical diagnosis challenging considering that some of the symptoms overlap with other vestibular disorders such vestibular migraine (VM) or autoimmune inner ear disease (AIED) (Hietikko et al. 2011; Lempert et al. 2012; Requena, Espinosa-Sanchez, and Lopez-Escamez 2014; Mijovic, Zeitouni, and Colmegna 2013).

Epidemiological evidence showing a genetic contribution in MD is based on familial aggregation studies with a high siblings recurrence risk ratio (λs= 16-48) (T. Requena et al. 2014) and the description of multiple familial cases in European and Asian descendant populations (Hietikko et al. 2013; Arweiler-Harbeck et al. 2011). Exome sequencing has identified private variants in the *FAM136A, DTNA, PRKCB, SEMA3D* and *DPT* genes in 4 families with autosomal dominant MD, showing incomplete penetrance and variable expressivity (Requena et al. 2015; Martín-Sierra et al. 2016, 2017). Moreover, some relatives in familial MD show partial syndromes, either with SNHL or episodic vertigo, increasing the granularity in the phenotype in a given family. However, the genetic contribution of familial genes in sporadic cases has not been investigated and the occurrence of recessive and de novo variation (DNV) is not known. More than 110 genes and ≈ 6000 variants have been related to hereditary non-syndromic hearing loss, making gene sequencing panels an essential tool for genetic diagnosis of hearing loss (Sloan-Heggen et al. 2016; The Molecular Otolaryngology and Renal Research Laboratories, The University of Iowa). Those genes include, 25 associated to autosomal dominant SNHL while 70 genes were related to recessive hearing loss (Shearer, Hildebrand, and Smith 1993).

Targeted gene sequencing panels have been demonstrated to be an excellent tool for molecular diagnosis of rare variants in known genes with allelic heterogeneity (Lionel et al. 2018; Brownstein et al. 2011), as well as in sporadic cases of hearing impairment in specific populations (Dallol et al. 2016; Gu et al. 2015). So, we selected SNHL genes and designed a custom gene panel to search for rare and DNV in sporadic MD.

In the present study, we describe the genetic variation found in a custom exon-sequencing panel of 69 SNHL genes in a large cohort of MD sporadic cases. We report that certain genes such as *GJB2*, *USH1G*, *SLC26A4* and *CLDN14* show an excess of missense variants in sporadic MD cases when compared to controls in the Iberian population, suggesting that several rare variants in these genes may contribute to the SNHL phenotype in sporadic MD.

## Materials and methods

### Editorial Policies and Ethical Considerations

This study protocol was approved by the Institutional Review Board for Clinical Research (MS/2014/02), and a written informed consent to donate biological samples was obtained from all subjects.

### Sample selection

A total of 890 Spanish and Portuguese patients with MD were recruited. All patients were diagnosed by neurotology experienced clinicians from the Meniere’s Disease Consortium (MeDiC), according to the diagnostic criteria for MD formulated by the International Classification Committee for Vestibular Disorders of the Barany Society in 2015 (Lopez-Escamez et al. 2015). Among them, 830 were considered sporadic MD cases while 60 were familial MD cases. Details from the selected cases are described in Table 1. As controls, 40 healthy individuals were selected from the same population.

**Table 1.** Participant individuals in this study. Number of individuals and geographical distribution of the selected cases and controls for targeted-gene sequencing. SMD, sporadic Meniere disease; FMD, familial Meniere disease.

| **Diagnosed** | *N* | **Sex** | % | **Location** | *N* |
| --- | --- | --- | --- | --- | --- |
| **SMD** | 830 | **Male** | 40 | **North** | 500 |
|  |  |  |  | **South** | 320 |
| **FMD** | 60 | **Female** | 60 | **Centre** | 90 |
| **Control** | 40 |  |  | **Portugal** | 20 |

### Selection of target genes

Target genes were selected from a literature search attending to human phenotype (hearing profile, comorbid vestibular symptoms) and phenotype observations in mouse and zebrafish models. Most of them were selected from HearingLoss.org website gene list for monogenic SNHL. Additional genes were added because they have been previously found in familial MD (Requena et al. 2015; Martín-Sierra et al. 2016, 2017), or allelic variations associated with hearing outcome in MD had been described, such as NFKB1 or TLR10 genes (Requena et al. 2013; Cabrera et al. 2014). Mitochondrial genes were added since maternal inheritance is suspected in several families with MD. Relevant information about the location, size, bibliography and other characteristics about each gene included in the panel is presented in Table S1 and Table S2.

The custom panel (Panel ID: 39351-1430751809) were designed by the Suredesign webtool (Agilent) to cover the exons and 50 bp in the flanking regions (5’ and 3’ UTR). This allowed the sequencing around 533.380kb with more than 98.46% coverage.

### Sample pooling

Enrichment technology allows the selective amplification of targeted sequences and DNA sample pooling, reducing the costs of reagents and increasing sample size. We decided to pool patient samples according to their geographical origin. Each pool consisted of 10 DNA samples from the same hospital for a total of 93 pools (930 samples).

DNA concentration and quality were measured on each sample using two methods: Qubit dsDNA BR Assay kit (TermoFisher Scientific) and Nanodrop 2000C (ThermoFisher Scientific). All samples had quality ratios ranging 1.8 and 2.0 in 280/260 and 1.6 to 2.0 in 260/230.

### Libraries preparation protocol

The HaloPlex Target Enrichment System (Agilent, Santa Clara, CA) was used to prepare the DNA libraries, according to the manufacturer protocol. Validation of the protocol and library performance was analyzed with a 2100 Bioanalyzer High Sensitivity DNA Assay kit. Expected concentrations were between 1 and 10 ng/ul. Higher concentrations than 10 ng/ul were diluted 1:10 in 10mM TRIS, 1mM EDTA. Targeted-sequencing was performed in an Illumina HiSeq 2000 platform.

### Data generation pipelines

Raw data downloaded and sequencing adapters were trimmed following manufacturer indications. The minimum coverage considered was 30X mean depth, however mitochondrial sequences reached higher coverages due to their shorter sizes. Bioinformatic analyses were performed according to the Good Practices recommended by Genome Analysis ToolKit (https://software.broadinstitute.org/gatk/).

Two methods were used to find differences in how UnifiedGenotyper and HaplotypeCaller (the old and the most recent tools for variant calling in GATK suite) address sequenced pools. Both custom pipelines use BWA-mem aligner and GATK suite tools following the GATK protocol for Variant Calling against GRCh37/hg19 human reference genome. Left normalization for multi-allelic variants were addressed by separated. Calling was made in the first pipeline with UnifiedGenotyper modifying number of chromosomes per sample (per pool, there are 20 chromosomes). The second pipeline used HaplotypeCaller, which cannot allow the same approach, but can automatically address high number of calls with a different approach. Variants with read depth (RD) <10 and genotype quality (GQ)<20 were excluded in all the calling pipelines following recommended hard filtering steps by GATK suite.

A third caller tool, VarScan, was used to filter and annotate quality strand data per variant to compare its output with GATK-based callers. VarScan allows the variant filtering using the information obtained according to each strand polarity. The method retrieves those variants that were only called in one strand, but not in the reverse strand, leading to false positive calls. This step was used as internal quality control to avoid strand bias usually generated in Haloplex data, as it has been reported in other studies (Collet et al. 2015).

### Positive control SNV validation

Positive control testing was addressed using samples from patients with familial MD with known variants on certain genes. These individuals come from previous familial studies with independently validated variants by Sanger sequencing. Known variants were also sequenced and validated by Sanger (Table S3). Coverage and mapping quality after each pipeline were annotated and measured. Representative chromatographs from validated SNVs are detailed in Figure S1.

### Selection and prioritization of pathogenic SNV

In order to obtain more information of each SNV, we annotated the merged files using the ANNOVAR tool. Minor allele frequencies (MAF) were retrieved for each candidate variant from gnomAD and ExAC database (total individuals and non-Finnish European (NFE) individuals). Since the estimated prevalence of sporadic MD in Spain is 0.75/1000 individuals (Morales Angulo et al. 2003), we selected variants with MAF <0.001 for single rare variant analysis. For burden analysis of common and rare variants, we chose a higher MAF value <0.1. The Collaborative Spanish Variant Server (CVCS) database including 1644 unrelated individuals was also used for annotation of exonic rare variants and to retrieve MAF in Spanish population (dataset fully accessible from http://csvs.babelomics.org/) (Dopazo et al. 2016).

KGGseq suite (grass.cgs.hku.hk/limx/kggseq) was used for the selection of rare variants and DNV to prioritize the most pathogenic variants according to the integrated model trained algorithm with known pathogenic variants and neutral control variants.

Enrichment analysis for each gene was made with all the exonic variants found with a MAF <0.1. This analysis required to divide the total amount of variants into three groups: those described in global gnomAD population, those found in NFE population, and finally those included in CVCS Spanish population. These three reference datasets were used for enrichment analysis comparison.

### Validation of candidate pathogenic SNV

Candidate SNV were visually revised in the BAM files using Integrative Genomics Viewer (https://software.broadinstitute.org/software/igv/) and validated in the different pools where they were called using Sanger sequencing.

### Population statistics

Statistical analysis was performed with IBM SPSS v.20 program, Microsoft Excel suite tools, and diverse python and java encoded public scripts. Due to the overrepresentation of Spanish population in our dataset, most of the selected variants were filtered through exome sequencing data from Spanish controls of CSVS database. The MAF was calculated for each variant in our dataset and rare and DNV on MD patients were identified in our gene panel. Odd ratios with 95% confidence interval were calculated for each variant using MAF obtained from Spanish population (N=1579), ExAC (N=60706), and ExAC NFE (N=33370) populations as controls.

Gene burden analysis was addressed using 2×2 contingency tables counting total exonic alternate allele counts per gene in our cases against total and NFE controls in ExAC and CSVS controls. Odds ratios with 95% confidence intervals were calculated using Fisher’s exact test and obtaining one-sided p-values. P-values were also corrected for multiple testing by the total amount of variants found for each gene following Bonferroni approach.

### Position of variants in significant enriched genes

Several models were generated for rare variant-enriched domains in significant enriched genes by using the INSIDER modelling tool (Meyer et al. 2018). The selected variants per gene are detailed in results. Prediction values were annotated with their calculated p values.

## Results

### Single rare variant analysis

A total of 2770 SNV were selected from the raw merged dataset (18961 SNVs) after filtering by quality controls. For rare variants analysis, SNV that were found in more than one pool were selected, remaining 1239 variants. After that, we filtered by variants observed in the control pools, leaving only 392 exonic SNV in cases (278 missense, 111 synonymous, 2 stopgain and 1 stoploss).

A final set of 162 SNV with a MAF <0.001 were retrieved (143 missense, 18 synonymous, 1 stoploss, 1 stopgain). All the exonic variants were annotated and scored using different priorization tools. Of them, 136 SNVs were not previously described in any population database and we considered them as possible de novo variants (DNVs).

After prioritizing the exonic variants, 31 rare variants remained (Table 2). Six of them were validated by Sanger sequencing in more than 2 individuals in the following genes: *GJB2*, *ESRRB*, *USH1G*, *SLC26A4* (Table S3). The rest of the variants were considered benign or likely benign since they did not reach the pathogenicity threshold predicted for KGGSeq. However, a novel synonymous variant in the *MARVELD2* gene was found and validated in 3 unrelated individuals.

**Table 2.**
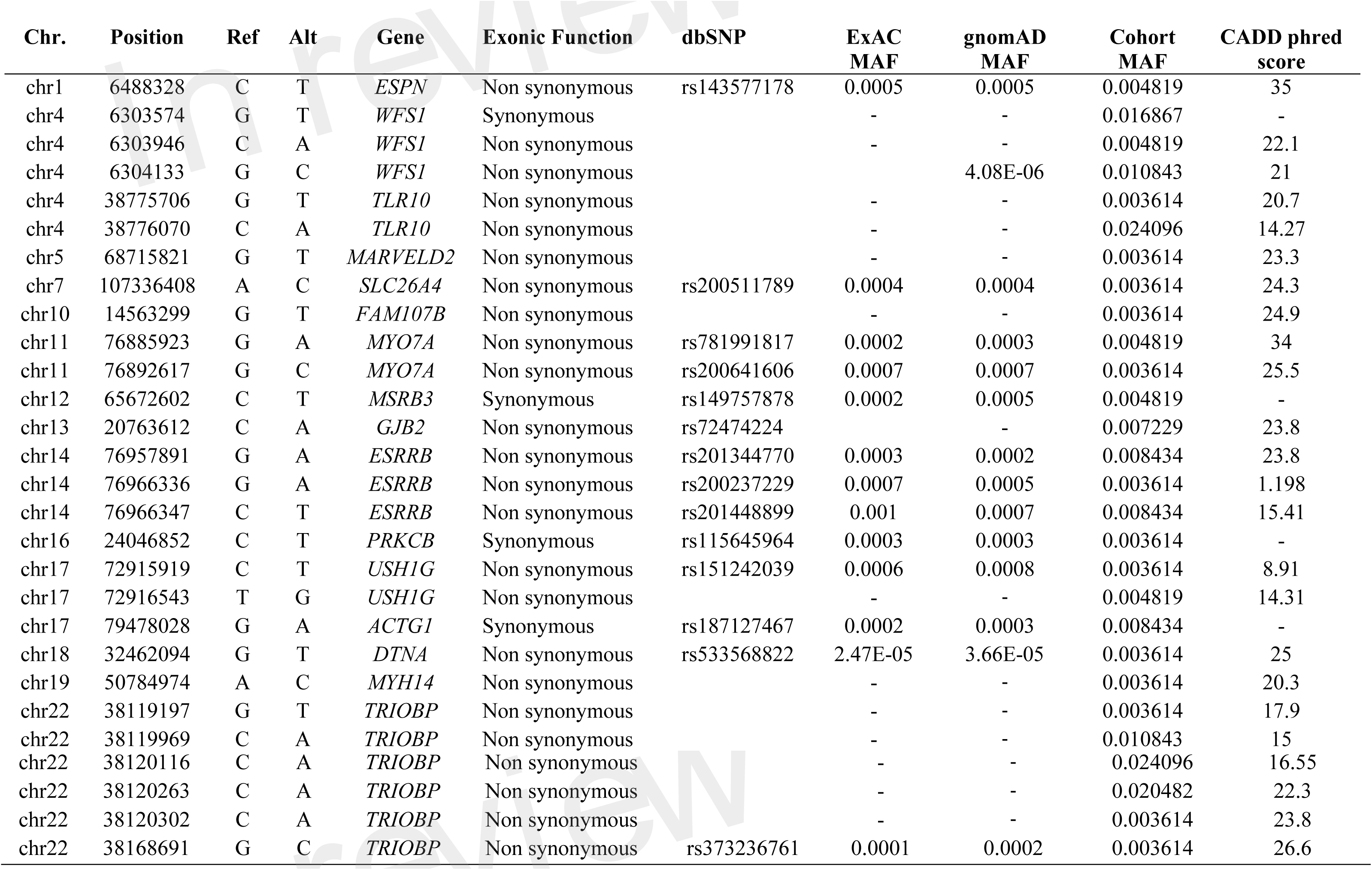
Prioritized rare SNVs found in the single rare variant analysis for sporadic MD cases. Minor allele frequency for each SNV is detailed as annotated by ExAC and gnomAD (exomes). Pathogenicity prediction is detailed according to CADD phred score.

The minor allelic frequencies in SNV of the 24 mitochondrial genes included in the panel were compared with the reference data obtained from MITOmap through its automated mtDNA sequence analysis system Mitomaster (Ruiz-Pesini et al. 2007). However, we did not found any SNV associated with MD (data not shown).

### Gene burden analysis

To analyse the interaction of multiple variants, we considered SNV with a MAF<0.1 for the gene burden analysis. A total of 957 exonic variants were retrieved and their frequencies were compared with the global and NFE frequencies from ExAC, and with the Spanish population frequencies from CSVS.

A gene burden analysis using our gene set was performed using these three reference datasets. After Bonferroni correction, some genes showed a significant enrichment of rare variants in the three comparisons, making them candidate genes to be selected for a diagnosis panel for MD (Table 3). Moreover, 6 genes (*FAM136A, ADD1, SLC12A2, POU4F3, RDX* and *PRKCB*) presented some novel variants that were validated by Sanger, but they have not been described in global ExAC or CSVS datasets. Although these variants could not be sequenced in all the parents of these patients, we considered them as DNV.

**Table 3.** Gene burden analysis 1. List of 29 genes showing a significant excess of missense exonic variants in patients with sporadic MD, according to the MAF observed in global ExAC population (N=60706), non-Finnish European ExAC population (NFE) (N=33370) and Spanish population from CSVS (N=1579).

| Gene | # variants | Odds ratio<br>ExAC<sup>a</sup> | P value | P corrected<sup>b</sup> | Odds ratio<br>ExAC NFE<sup>a</sup> | P value | P corrected<sup>b</sup> | Odds ratio<br>CSVS<sup>a</sup> | P value | P corrected<sup>b</sup> |
| --- | --- | --- | --- | --- | --- | --- | --- | --- | --- | --- |
| MYH14 | 50 | 13,54 (5,85-31,37) | 1.20E-09 | 6.01E-08 | 56,94 (25,13-128,98) | <1.00E-15 | <1.00E-15 | 38,52 (16,95-87,56) | <1.00E-15 | <1.00E-15 |
| MYO7A | 43 | 3,11 (1,73-5,57) | 1.41E-04 | 6.08E-03 | 4,64 (2,65-8,13) | 7.72E-08 | 3.32E-06 | 3,85 (2,18-6,81) | 3.60E-06 | 1.55E-04 |
| ADD1 | 29 | 249,89 (96,17-649,29) | <1.00E-15 | <1.00E-15 | 318,03 (122,45-826) | <1.00E-15 | <1.00E-15 | 747,23 (287,95-1939,08) | <1.00E-15 | <1.00E-15 |
| WHRN | 28 | 24,63 (7,96-76,17) | 2.68E-08 | 7.51E-07 | 39,97 (13,03-122,58) | 1.12E-10 | 3.13E-09 | 62,91 (20,62-191,98) | 3.44E-13 | 9.64E-12 |
| TPRN | 26 | 36,63 (10,61-126,49) | 1.24E-08 | 3.22E-07 | 46,93 (13,64-161,49) | 1.03E-09 | 2.68E-08 | 81,9 (23,93-280,26) | 2.24E-12 | 5.81E-11 |
| USH1G | 24 | 4,04 (1,91-8,51) | 2.46E-04 | 5.90E-03 | 18,38 (9,26-36,48) | <1.00E-15 | <1.00E-15 | 8,12 (4-16,49) | 6.57E-09 | 1.58E-07 |
| USH1C | 22 | 5,12 (2,58-10,17) | 3.13E-06 | 6.88E-05 | 10,68 (5,54-20,59) | 1.54E-12 | 3.38E-11 | 5,12 (2,58-10,18) | 3.06E-06 | 6.73E-05 |
| P2RX2 | 22 | 20,34 (9,75-42,44) | <1.00E-15 | <1.00E-15 | 23,74 (11,41-49,41) | <1.00E-15 | <1.00E-15 | 23,51 (11,29-48,94) | <1.00E-15 | <1.00E-15 |
| ESPN | 19 | 27,88 (17,06-45,55) | <1.00E-15 | <1.00E-15 | 24,35 (14,88-39,84) | <1.00E-15 | <1.00E-15 | 17,39 (10,59-28,56) | <1.00E-15 | <1.00E-15 |
| RDX | 17 | 94,37 (36,06-247) | <1.00E-15 | <1.00E-15 | 57,38 (21,85-150,66) | <1.00E-15 | <1.00E-15 | 75,26 (28,72-197,23) | <1.00E-15 | <1.00E-15 |
| SLC26A4 | 15 | 1,79 (1,02-3,15) | 4.31E-02 | 6.46E-01 | 4,14 (2,5-6,86) | 3.37E-08 | 5.06E-07 | 3,65 (2,19-6,09) | 6.62E-07 | 9.93E-06 |
| ESRRB | 14 | 12,54 (7,26-21,63) | <1.00E-15 | <1.00E-15 | 10,67 (6,16-18,48) | <1.00E-15 | <1.00E-15 | 7,54 (4,31-13,18) | 1.41E-12 | 1.98E-11 |
| DTNA | 13 | 6,07 (4,68-7,87) | <1.00E-15 | <1.00E-15 | 4,87 (3,74-6,35) | <1.00E-15 | <1.00E-15 | 4,78 (3,67-6,23) | <1.00E-15 | <1.00E-15 |
| PRKCB | 13 | 27,62 (18,65-40,91) | <1.00E-15 | <1.00E-15 | 17,6 (11,84-26,16) | <1.00E-15 | <1.00E-15 | 40,79 (27,61-60,28) | <1.00E-15 | <1.00E-15 |
| SLC12A2 | 12 | 21,91 (15,95-30,1) | <1.00E-15 | <1.00E-15 | 55,14 (40,31-75,43) | <1.00E-15 | <1.00E-15 | 79,32 (58,04-108,41) | <1.00E-15 | <1.00E-15 |
| KCNQ1 | 11 | 7,93 (5,92-10,63) | <1.00E-15 | <1.00E-15 | 13,75 (10,33-18,29) | <1.00E-15 | <1.00E-15 | 6,96 (5,18-9,34) | <1.00E-15 | <1.00E-15 |
| NFKB1 | 10 | 1,87 (1,3-2,71) | 7.81E-04 | 7.81E-03 | 1,92 (1,33-2,77) | 4.69E-04 | 4.69E-03 | 3,79 (2,72-5,29) | 4.44E-15 | 4.44E-14 |
| **DPT** | 10 | 1,88 (1,2-2,92) | 5.36E-03 | 5.36E-02 | 8,03 (5,5-11,74) | <1.00E-15 | <1.00E-15 | 4,38 (2,94-6,51) | 3.01E-13 | 3.01E-12 |
| **CEACAM16** | 9 | 1,25 (1,04-1,49) | 1.56E-02 | 1.40E-01 | 1,33 (1,12-1,59) | 1.48E-03 | 1.33E-02 | 2,45 (2,09-2,87) | <1.00E-15 | <1.00E-15 |
| **FAM136A** | 8 | 31,18 (27,22-35,71) | <1.00E-15 | <1.00E-15 | 38,36 (33,5-43,92) | <1.00E-15 | <1.00E-15 | 14,71 (12,81-16,89) | <1.00E-15 | <1.00E-15 |
| **GRHL2** | 8 | 11,72 (9,35-14,7) | <1.00E-15 | <1.00E-15 | 9,28 (7,39-11,66) | <1.00E-15 | <1.00E-15 | 6,45 (5,11-8,15) | <1.00E-15 | <1.00E-15 |
| **EYA4** | 7 | 36,65 (28,83-46,59) | <1.00E-15 | <1.00E-15 | 25,31 (19,88-32,22) | <1.00E-15 | <1.00E-15 | 7,68 (5,97-9,88) | <1.00E-15 | <1.00E-15 |
| **COCH** | 7 | 3,41 (2,54-4,58) | <1.00E-15 | <1.00E-15 | 3,39 (2,52-4,56) | <1.00E-15 | <1.00E-15 | 3,95 (2,95-5,28) | <1.00E-15 | <1.00E-15 |
| **CCDC50** | 5 | 1,25 (0,99-1,59) | 6.22E-02 | 3.11E-01 | 1,62 (1,3-2,03) | 2.51E-05 | 1.25E-04 | 3,22 (2,63-3,94) | <1.00E-15 | <1.00E-15 |
| **KCNJ10** | 5 | 4,38 (3,9-4,93) | <1.00E-15 | <1.00E-15 | 3,14 (2,78-3,55) | <1.00E-15 | <1.00E-15 | 2,3 (2,02-2,61) | <1.00E-15 | <1.00E-15 |
| **SEMA3D** | 4 | 1,56 (1,03-2,38) | 3.60E-02 | 1.44E-01 | 1,13 (0,72-1,77) | 5.84E-01 | 2.34E+00 | 3,79 (2,63-5,47) | 1.09E-12 | 4.35E-12 |
| **CLDN14** | 4 | 4,51 (2,56-7,94) | 1.74E-07 | 6.97E-07 | 22,86 (13,56-38,55) | <1.00E-15 | <1.00E-15 | 4,72 (2,69-8,29) | 6.61E-08 | 2.65E-07 |
| **POU4F3** | 4 | 7,31 (6,53-8,18) | <1.00E-15 | <1.00E-15 | 4,92 (4,38-5,52) | <1.00E-15 | <1.00E-15 | 9,2 (8,23-10,28) | <1.00E-15 | <1.00E-15 |
| **MSRB3** | 4 | 13,35 (11,34-15,71) | <1.00E-15 | <1.00E-15 | 10,85 (9,21-12,79) | <1.00E-15 | <1.00E-15 | 4,06 (3,4-4,83) | <1.00E-15 | <1.00E-15 |

<sup>a</sup>Odds ratios were calculated in the 95% confidence interval.
<sup>b</sup>P values were corrected with Bonferroni method approach.

A second variant analysis using the missense variants described in CSVS Spanish population database was made (Table 4). Eighteen genes showed an excess of missense variants (a total of 46 variants, detailed in Table S4). Of note, five genes causing autosomal recessive SNHL showed the highest accumulation of missense variants when they were compared with NFE and Spanish population datasets: *SLC26A4*, *GJB2, CLDN14, ESRRB* and *USH1G*. The variants in these five genes were validated through Sanger sequencing and considered Spanish population-specific variants.

**Table 4.**
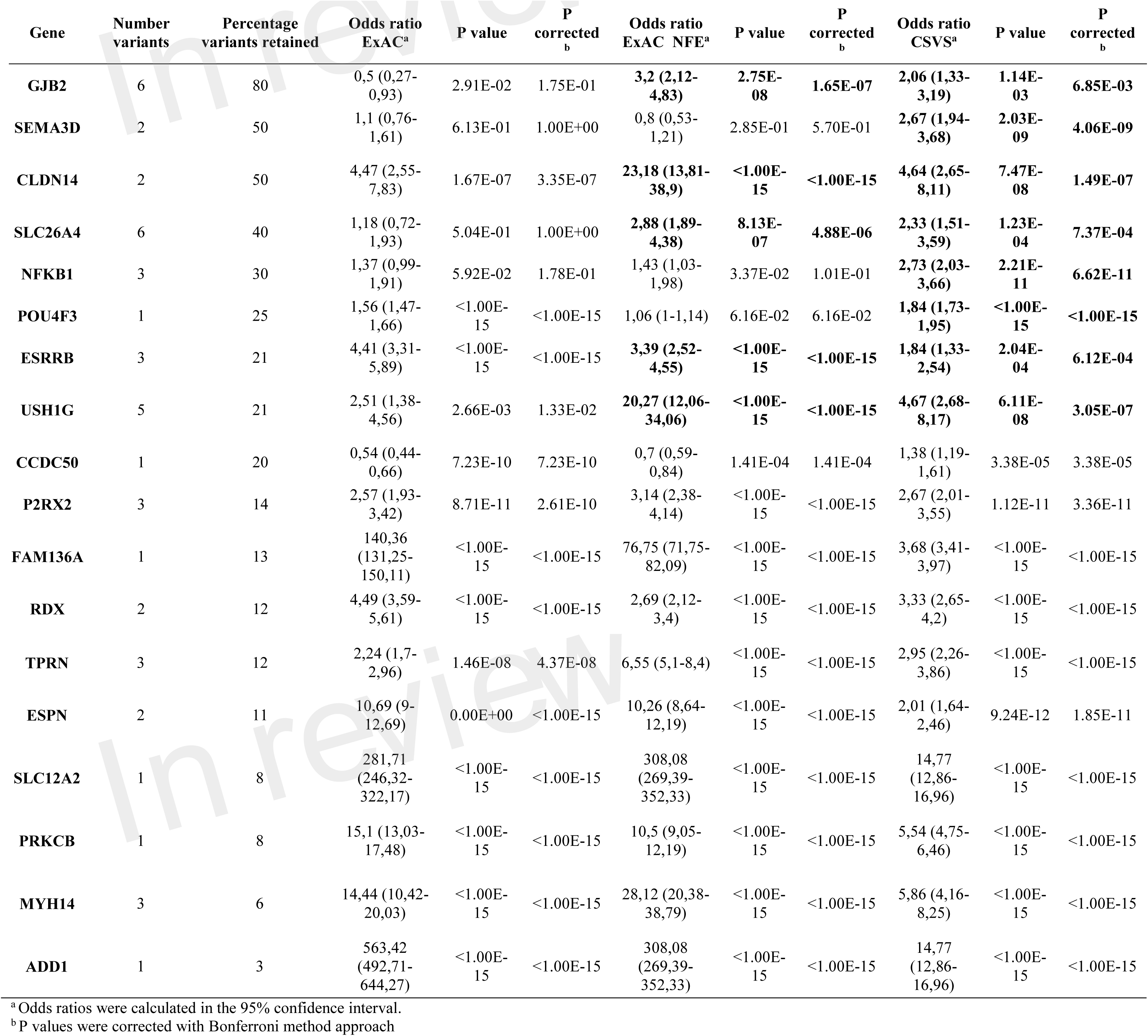
Gene burden analysis 2. List of 18 genes showing a significant excess of missense exonic variants in patients with sporadic MD, according to the MAF observed in CSVS Spanish database (N=1579), compared with global ExAC population (N=60706) and non-Finnish European ExAC population (N=33370). Selected SNV can be consulted in Table S4.

### Excess of rare variants in hearing loss genes in familial cases

We used exome sequencing datasets from familial MD cases previously reported to search for rare variants identified in our panel in the sporadic cases. Although no single missense variant was found segregated in all the cases in the same family, we found several rare missense variants in at least one case per family in genes such as *GJB2*, *GRHL2*, *TRIOBP*, *RDX*, *KCNQ4*, *WFS1* and *ADD1*. These MD families show phenotypic differences in terms of age of onset, hearing profile and disease progression and the presence of rare variants can be addressed as potential modulators of the phenotype in each familial case (Table 5).

**Table 5.**
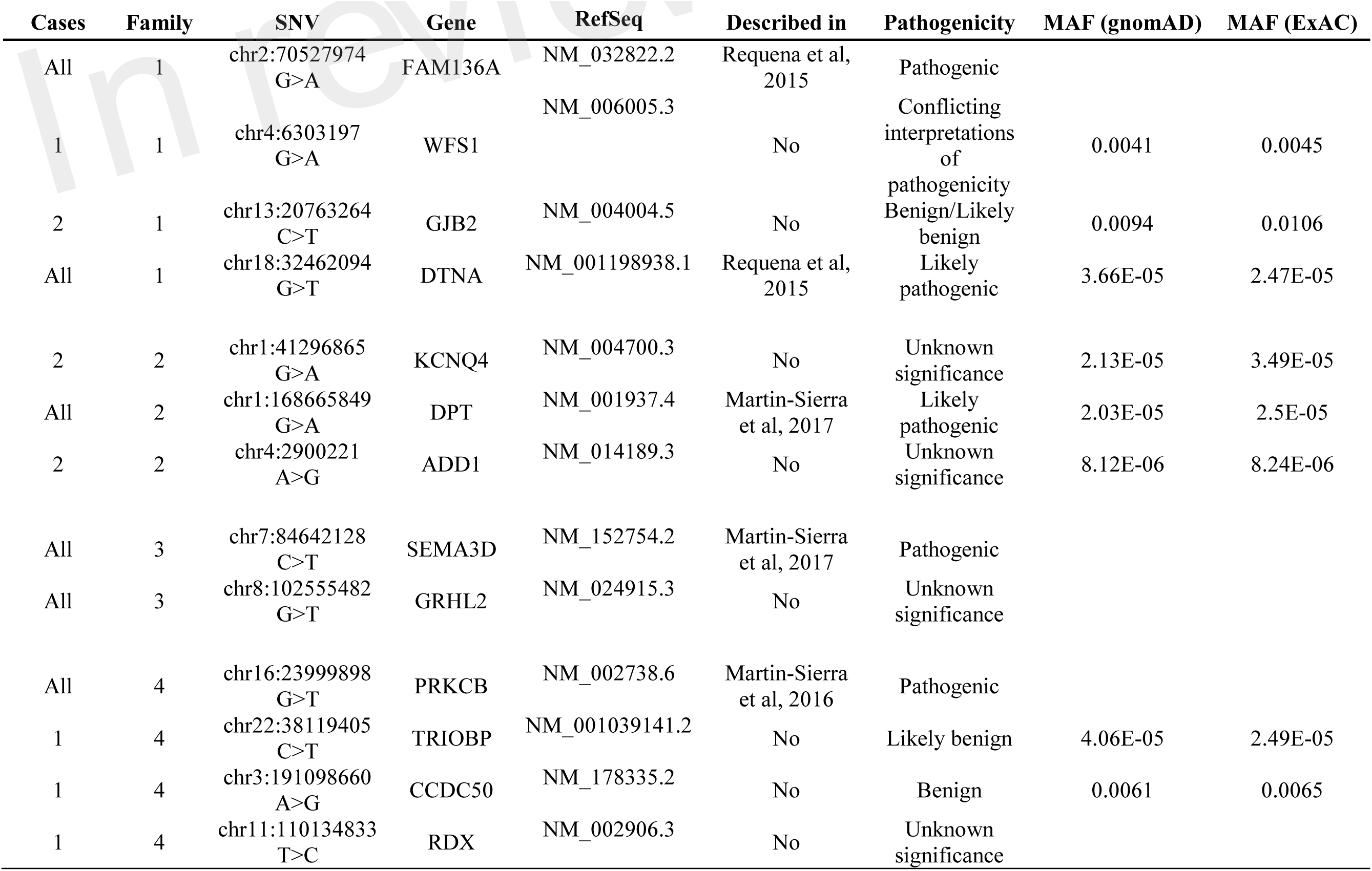
Missense variants found in familial MD cases. Variants were retrieved from familial cases segregating a partial phenotype in different families.

### Effect of rare variant interaction

We selected exonic variants from the gene burden analysis to analyze their potential additive effect at protein-protein interaction interfaces by the tool INSIDER for our selected five genes. However, protein interfaces for *ESRRB*, *CLDN14* and *SLC26A4* genes could not be loaded and processed on the database (lacking predicted interfaces on ÉCLAIR database or crystalized protein structures on Protein Data Bank (PDB) database). Of note, most relevant affected interaction is observed in the self-interaction *GJB2*-*GJB2* by the known variants observed in the burden analysis (significant spatial clustering with 4 SNV, p=0.0009) rs111033218:G>C, rs80338945:A>G, rs374625633:T>C and rs2274084:C>T (Fig 1A).

**Figure 1.**
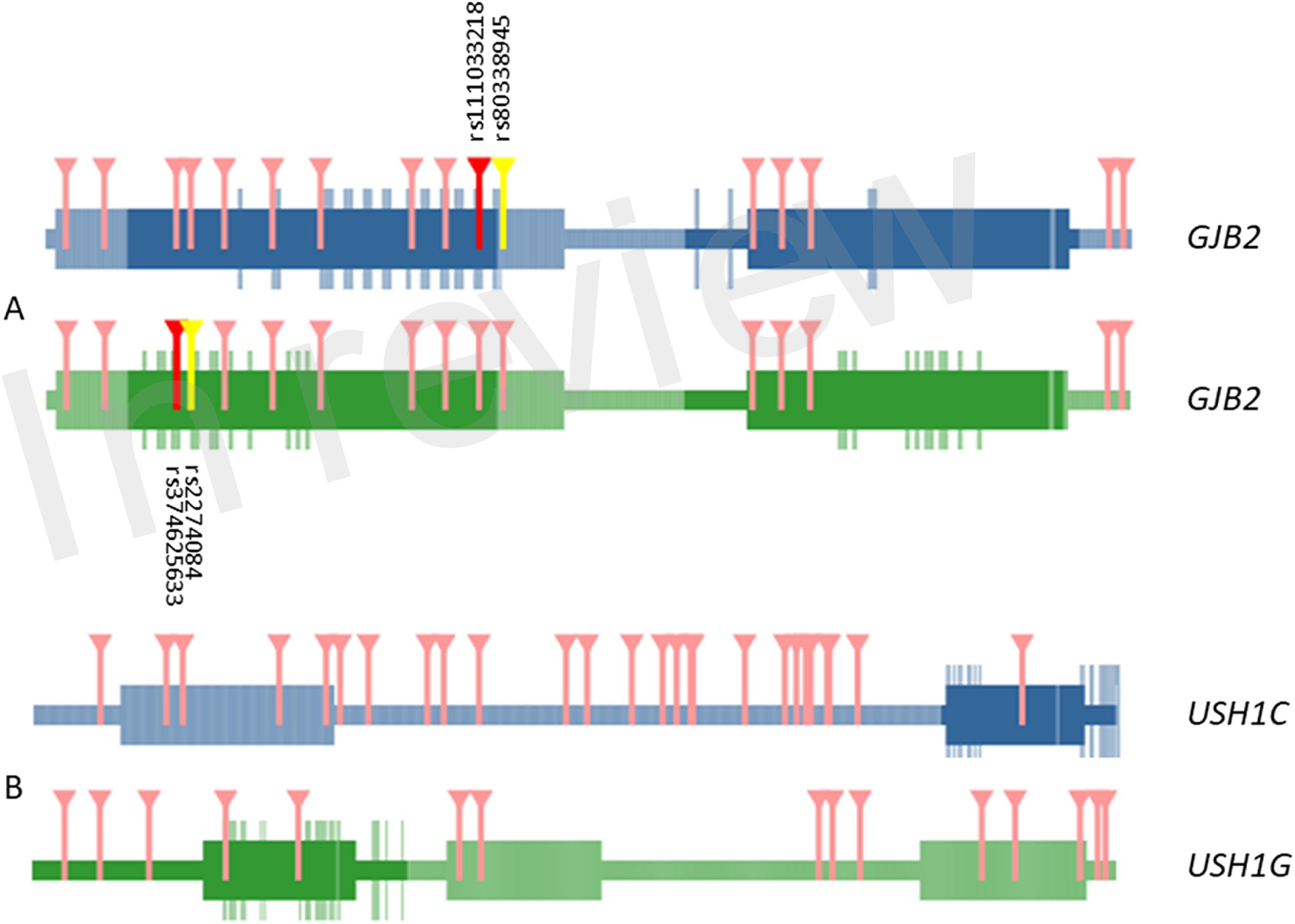
Representation of domains and interactive interface in GJB2 – GJB2 (A) and USH1G – USH1C (B) interaction. Marked in darker color boxes, functional regions of the protein. Aminoacids in the interactive surface of the protein are highlighted in the same color. Variants that affect the interaction regions between both proteins are marked in red and yellow. Only in GJB2 – GJB2 self-interaction missense mutations are relevant in the interaction (dbSNP ids detailed in black). The rest of the variants affecting aminoacids tested in both interactions that are out in the interactive surface region are marked in pink.

Other interactions of interest were founded between the *USH1G* – *USH1C* genes, but the involved variants were not located in the known interaction surface of *USH1G* (Fig 1B).

## Discussion

This study shows that patients with sporadic MD have an enrichment of few rare variants in certain hearing loss genes such as *GJB2*, *SLC26A* or *USH1G*. This excess of missense variants in some genes may increase the risk to develop hearing loss in MD and may contribute to explain the heterogeneity observed in the phenotype (Francioli et al. 2015). To understand the relevance of population frequencies in our cohort, we performed the association analysis between variants observed in MD cases against their respective frequencies on a healthy population for each gene of the panel. From the total amount of variants, we selected exonic variants for all the targeted genes. We applied a stronger filter for the selection of missense variants by choosing previously described variants for each gene significantly overrepresented in MD cases.

Since many missense variants were not found in the Spanish population from CSVS, a third comparison limited to those variants described in CSVS database was carried out. We followed this conservative approach to reduce false positive findings in the readings. This third restrictive gene analysis was limited to 132 variants observed at least once in the Spanish reference population.

From the final analysis, we found that some genes such as *SLC26A4*, *ESRRB*, *CLDN14*, *GJB2* and *USH1G* retained the higher number of missense variants among Spanish MD patients. We also found one synonymous variant in the *MARVELD2* gene. Besides from its functional implications, it may also generate a cryptic splice site. However, more testing is needed to confirm this hypothesis.

### Gene panel for familial MD

The Genomics England project (https://www.genomicsengland.co.uk/) has designed gene panels for the diagnosis of many genetic disorders including familial MD (https://panelapp.genomicsengland.co.uk/panels/394/). This panel is in an early stage of development because it only considers 130 genes with limited evidence to few families. The results of this study can be used to improve the design of panels for the diagnosis of MD.

For the design of our panel, we chose a total of 69 genes. Most of the genes were selected according to the hearing loss profile (low frequency or pantonal hearing loss). However, more than 90 genes have been related to hearing loss, so more hearing loss genes could be involved in the phenotype (fully accessible from Hereditary Hearing Loss Homepage: http://hereditaryhearingloss.org/). Genetic evidence of hearing loss has been obtained from linkage analyses until the emergence of NGS techniques (Shearer, Hildebrand, and Smith 1993), that have facilitated the clinical development of genetic diagnosis in hearing loss. Custom panels and microarrays have been the flags of a new age of discovery of novel and rare variants for genetic diagnostic of hearing loss (Brownstein et al. 2011; Shearer and Smith 2015).

Our panel was designed considering hearing loss as the main symptom shared by all patients with MD, since the vestibular phenotype and other associated co-morbidities such as migraine or autoimmune disorders are more variable. To improve the diagnostic yield of MD and to decrease this granularity in the phenotype, it will be recommendable to select sporadic patients with an early age of onset for future studies.

### Rare variants in hearing loss genes in sporadic MD

The frequency of hearing loss related genes is population-specific (Sloan-Heggen et al. 2016). Herein, we present a study for MD patients in the Spanish population. As a part of the study, we consider a panel of genes related to hearing loss and other symptoms. Besides from the validated variants in singletons, only a few rare variants such as *ESRRB* rs201448899:C>T, *MARVELD2* rs369265136:G>A, *SLC26A4* rs200511789:A>C and *USH1G* rs151242039:C>T have been validated in more than one sporadic case in the entire cohort. All these genes had been previously considered as pathogenic for hearing loss, but they have never been involved with MD.

*ESRRB* is the estrogen-related receptor beta, also known as nuclear receptor subfamily 3, group B, member 2 or NR3B2. This gene encodes for a protein like the estrogen receptor but with a different and unknown role. Mutations in the mouse orthologue have been involved in the placental development and autosomal recessive SNHL (Collin et al. 2008; Weber et al. 2014).

*MARVELD2* encodes a protein found in the tight junctions, between epithelial cells. The encoded protein seems to forge barriers between epithelial cells such the ones in the organ of Corti, Defects in this gene are associated with DFNB49 (Mašindová et al. 2015).

*SLC26A4* gene encodes pendrin, a protein extensively studied in hearing loss. Its alteration is one of the most common causes of syndromic deafness and autosomal recessive SNHL. It is also associated with enlarged vestibular aqueduct syndrome (EVAS) (Yang et al. 2007, 36).

*USH1G* is a gene translating to a protein that contains three ankyrin domains, a class I PDZ-binding motif and a sterile alpha motif. This protein is well-known to interact with harmonin (*USH1C*) in the stereocilia of hair cells, a protein associated with Usher syndrome type 1C (Weil et al. 2003). This protein plays a role in the development and maintenance of the auditory and visual systems and functions in the cohesion of hair bundles formed by inner ear sensory cells. Alterations in the integrity of the protein seem to be the cause of Usher syndrome type 1G (Miyasaka et al. 2016; Weil et al. 2003).

However, *ESRRB* rs201448899:C>T has been observed in more Spanish controls than in global or NFE in ExAC. This increased frequency on the Iberian population when compared with other known largest frequencies as NFE, suggests that this is a population specific variant rather than a MD disease variant. Only the *MARVELD2* rs369265136:G>A variant remains as a proper DNV related to MD cases. However, the functional effect of a synonymous variant is unknown and functional studies will be required to decipher the relevancy of this variant in MD cases in the future.

### Burden analysis of rare missense variants in sporadic MD

Our results demonstrate a burden of rare missense variants in few SNHL genes, including *GJB2, ESRRB, CLDN14, SLC26A4* and *USH1G*. We speculate that the additive effect of several missense variants in the same gene could interact with the same or other genes at the protein level resulting in the hearing loss phenotype.

Population analysis was addressed in order to obtain a better image of our cohort. Despite the limitation that represents the small number of genes considered in our panel, we have found a significant increase of missense variants on several hearing loss genes in the Iberian population (Table 4). These findings suggest the involvement of multiple missense variants in the same gene and may explain several clinical findings in MD. So, incomplete phenotype found in relatives of patients with familial MD or even the variable expressivity observed could be explained on the differences found in multiple rare variants with additive effect among individuals of the same family (Requena et al. 2015; Martín-Sierra et al. 2016, 2017). In addition, some sporadic cases where a single rare variant with unknown significance cannot explain the phenotype could be singletons individuals with several low frequency variants probably following a compound heterozygous recessive pattern of inheritance. Our results start to decipher the complex interaction between rare and ultrarare variations (MAF< 0.0001) in the same or different genes in sporadic MD, adding more evidence to understand the genetic architecture of MD.

First, the phenotype of MD could be the result of additive effect of low frequency or rare variants in the same gene. A good number of common or low frequency variants in the same gene can be a rare situation, as rare as the disease. As much changes are added to the protein, its integrity could be affected, showing a suboptimal functioning and finally, a loss of function. In our case, GJB2, that forms a hexamer with a transmembrane channel function, has been determined as possible affected by these changes in their interactions. Previous studies have determined how certain changes in the monomer can affect to the develop of the hexamer hemichannel (Bicego et al. 2006; Jara et al. 2012). Here, bioinformatics models show how the interaction of low frequency variants found in MD patients can impact the interaction between two connexins monomers, but this effect could be amplified in a model including the 6 connexins that form the connexon.

A second hypothesis points to the interaction of several genes in the disease phenotype, following its complex disease definition (Becker 2004; Mitchell 2012). In this case, high significant genes in our study could be added to the panel of candidate targets of the disease, although a single candidate variant could not be enough to explain the disease. So, the interaction between some of our selected genes and other, a priori, not related SNHL genes could be relevant in the expressivity. *USH1G* interacts with *USH1C*, a known gene involved in Usher syndrome. *USH1G* has been observed to have a minor role in Usher syndrome in Spanish population (Aller et al. 2007), but not in MD, even though they share similar hearing loss profile. Although no one of the missense variants in *USH1G* were in an interaction domain, this could be of interest when considering interaction between different proteins as a main factor to develop a mild phenotype. This hypothesis was reinforced through the data found in familial cases. For instance, the variant rs748718975 in *DPT* gene was only associated with the SNHL phenotype in the family where it was described, but these cases showed different characteristics in the age of onset or hearing loss outcome. These differences between the cases can be explained with other variants found in *KCNQ4* (rs574794136:G>A) and *ADD1* (rs372777117:A>G) genes, although these variants were previously described as variants of unknown significant. So, the variant rs574794136:G>A was found in two sisters with MD, but not in the third one, that was carrier of rs372777117:A>G. This excess of rare variants in certain genes observed in familial cases could explain the differences in expressivity in a given family.

This panel was made as an early screening diagnostic panel. Here we have found that certain SNHL gene variants can be related to MD in the Iberian population and the results show that multiple rare allelic variants in the same gene should be consider as likely pathogenic. Our results will contribute to design a better gene panel for the genetic diagnosis of MD.

## Author Contributions Statement

TR and JALE as project managers conceived the main idea of the work. Sample preparation and protocol were carry out by TR and AGM. AGM performed the bioinformatics and statistical analysis. Validations of tested SNV were done by AGM and PRN. AGM and JALE took the lead in writing the manuscript. All authors provided critical feedback and helped shape the research, analysis and manuscript.

## Data Availability Statement

The raw data supporting the conclusions of this manuscript will be made available by the authors, without undue reservation, to any qualified researcher.

## Acknowledgements

We acknowledge to all members of the Meniere disease Consortium (MeDiC), a network of clinical and research centres contributing to the study of Meniere disease. List of participants in MeDiC: Juan Carlos Amor-Dorado (Hospital Can Misses Ibiza, Spain), Ismael Aran (Complexo Hospitalario de Pontevedra, Spain), Angel Batuecas-Caletrío (Hospital Universitario Salamanca, Spain), Jesus Benitez (Hospital Universitario de Gran Canaria Dr. Negrin, Las Palmas de Gran Canaria, Spain), Jesus Fraile (Hospital Miguel Servet, Zaragoza, Spain), Ana Garcia-Arumí (Hospital Universitario Vall d’Hebron, Barcelona, Spain), Rocio Gonzalez-A (Hospital Universitario Marqués de Valdecilla, Santander, Spain), Juan M Espinosa-Sanchez (Hospital Virgen de las Nieves, Granada, Spain), Raquel Manrique Huarte, Nicolas Perez-Fernandez (Clinica Universidad de Navarra, Spain), Pedro Marques (Centro Hospitalar de São João, Porto, Portugal), EM-S, Ricardo Sanz (Hospital Universitario de Getafe, Madrid, Spain), Manuel Oliva Dominguez (Hospital Costa del Sol Marbella, Spain), PP (Hospital Cabueñes, Asturias, Spain), HP-G, VP-G (Hospital La Fe, Valencia, Spain), SS-P, AS-V (Complexo Hospitalario Universitario, Santiago de Compostela, Spain), MT (Instituto Antolí Candela, Madrid, Spain), Roberto Teggi (San Raffaelle Scientific Institute, Milan, Italy), and GT (Complejo Hospitalario Badajoz, Spain). We also acknowledge Joaquin Dopazo (Clinical Bioinformatics Area, Hospital Virgen del Rocio, Sevilla, Spain) to facilitate us the access to CSVS datasets. AGM is a PhD student in the Biomedicine program at University of Granada and this work is part of his doctoral thesis.

## References

Aller, Elena, Teresa Jaijo, Magdalena Beneyto, Carmen Nájera, Constantino Morera, Herminio Pérez-Garrigues, Carmen Ayuso, and Jose Millán. 2007. “Screening of the USH1G Gene among Spanish Patients with Usher Syndrome. Lack of Mutations and Evidence of a Minor Role in the Pathogenesis of the Syndrome.” Ophthalmic Genetics 28 (3): 151–55. https://doi.org/10.1080/13816810701537374.

Arweiler-Harbeck, Diana, Bernhard Horsthemke, Klaus Jahnke, and Hans Christian Hennies. 2011. “Genetic Aspects of Familial Ménière’s Disease.” Otology & Neurotology: Official Publication of the American Otological Society, American Neurotology Society [and] European Academy of Otology and Neurotology 32 (4): 695–700. https://doi.org/10.1097/MAO.0b013e318216074a.

Becker, Kevin G. 2004. “The Common Variants/Multiple Disease Hypothesis of Common Complex Genetic Disorders.” Medical Hypotheses 62 (2): 309–17. https://doi.org/10.1016/S0306-9877(03)00332-3.

Bicego, Massimiliano, Martina Beltramello, Salvatore Melchionda, Massimo Carella, Valeria Piazza, Leopoldo Zelante, Feliksas F. Bukauskas, et al. 2006. “Pathogenetic Role of the Deafness-Related M34T Mutation of Cx26.” Human Molecular Genetics 15 (17): 2569–87. https://doi.org/10.1093/hmg/ddl184.

Brownstein, Zippora, Lilach M. Friedman, Hashem Shahin, Varda Oron-Karni, Nitzan Kol, Amal Abu Rayyan, Thomas Parzefall, et al. 2011. “Targeted Genomic Capture and Massively Parallel Sequencing to Identify Genes for Hereditary Hearing Loss in Middle Eastern Families.” Genome Biology 12 (9): R89. https://doi.org/10.1186/gb-2011-12-9-r89.

Cabrera, Sonia, Elena Sanchez, Teresa Requena, Manuel Martinez-Bueno, Jesus Benitez, Nicolas Perez, Gabriel Trinidad, et al. 2014. “Intronic Variants in the NFKB1 Gene May Influence Hearing Forecast in Patients with Unilateral Sensorineural Hearing Loss in Meniere’s Disease.” PloS One 9 (11): e112171. https://doi.org/10.1371/journal.pone.0112171.

Caulley, Lisa, Alexandra Quimby, Jacob Karsh, Azin Ahrari, Darren Tse, and Georgios Kontorinis. 2017. “Autoimmune Arthritis in Ménière’s Disease: A Systematic Review of the Literature.” Seminars in Arthritis and Rheumatism, November. https://doi.org/10.1016/j.semarthrit.2017.11.008.

Collet, Agnès, Julien Tarabeux, Elodie Girard, Catherine Dubois DEnghien, Lisa Golmard, Vivien Deshaies, Alban Lermine, et al. 2015. “Pros and Cons of HaloPlex Enrichment in Cancer Predisposition Genetic Diagnosis.” Genetics 2015, Vol. 2, Pages 263–280, December. https://doi.org/10.3934/genet.2015.4.263.

Collin, Rob W.J., Ersan Kalay, Muhammad Tariq, Theo Peters, Bert van der Zwaag, Hanka Venselaar, Jaap Oostrik, et al. 2008. “Mutations of ESRRB Encoding Estrogen-Related Receptor Beta Cause Autosomal-Recessive Nonsyndromic Hearing Impairment DFNB35.” American Journal of Human Genetics 82 (1): 125–38. https://doi.org/10.1016/j.ajhg.2007.09.008.

Dallol, Ashraf, Kamal Daghistani, Aisha Elaimi, Wissam A. Al-Wazani, Afaf Bamanie, Malek Safiah, Samira Sagaty, et al. 2016. “Utilization of Amplicon-Based Targeted Sequencing Panel for the Massively Parallel Sequencing of Sporadic Hearing Impairment Patients from Saudi Arabia.” BMC Medical Genetics 17 (Suppl 1). https://doi.org/10.1186/s12881-016-0329-8.

Dopazo, Joaquín, Alicia Amadoz, Marta Bleda, Luz Garcia-Alonso, Alejandro Alemán, Francisco García-García, Juan A. Rodriguez, et al. 2016. “267 Spanish Exomes Reveal Population-Specific Differences in Disease-Related Genetic Variation.” Molecular Biology and Evolution 33 (5): 1205–18. https://doi.org/10.1093/molbev/msw005.

Francioli, Laurent C., Paz P. Polak, Amnon Koren, Androniki Menelaou, Sung Chun, Ivo Renkens, Cornelia M. van Duijn, et al. 2015. “Genome-Wide Patterns and Properties of de Novo Mutations in Humans.” Nature Genetics 47 (7): 822–26. https://doi.org/10.1038/ng.3292.

Gazquez, Irene, Andres Soto-Varela, Ismael Aran, Sofia Santos, Angel Batuecas, Gabriel Trinidad, Herminio Perez-Garrigues, Carlos Gonzalez-Oller, Lourdes Acosta, and Jose A. Lopez-Escamez. 2011. “High Prevalence of Systemic Autoimmune Diseases in Patients with Menière’s Disease.” PloS One 6 (10): e26759. https://doi.org/10.1371/journal.pone.0026759.

Gu, X., L. Guo, H. Ji, S. Sun, R. Chai, L. Wang, and H. Li. 2015. “Genetic Testing for Sporadic Hearing Loss Using Targeted Massively Parallel Sequencing Identifies 10 Novel Mutations.” Clinical Genetics 87 (6): 588–93. https://doi.org/10.1111/cge.12431.

Hietikko, Elina, Jouko Kotimäki, Erna Kentala, Tuomas Klockars, Martti Sorri, and Minna Männikkö. 2011. “Finnish Familial Meniere Disease Is Not Linked to Chromosome 12p12.3, and Anticipation and Cosegregation with Migraine Are Not Common Findings.” Genetics in Medicine: Official Journal of the American College of Medical Genetics 13 (5): 415–20. https://doi.org/10.1097/GIM.0b013e3182091a41.

Hietikko, Elina, Jouko Kotimäki, Martti Sorri, and Minna Männikkö. 2013. “High Incidence of Meniere-like Symptoms in Relatives of Meniere Patients in the Areas of Oulu University Hospital and Kainuu Central Hospital in Finland.” European Journal of Medical Genetics 56 (6): 279–85. https://doi.org/10.1016/j.ejmg.2013.03.010.

Jara, Oscar, Rodrigo Acuña, Isaac E. García, Jaime Maripillán, Vania Figueroa, Juan C. Sáez, Raúl Araya-Secchi, et al. 2012. “Critical Role of the First Transmembrane Domain of Cx26 in Regulating Oligomerization and Function.” Molecular Biology of the Cell 23 (17): 3299–3311. https://doi.org/10.1091/mbc.E11-12-1058.

Lempert, Thomas, Jes Olesen, Joseph Furman, John Waterston, Barry Seemungal, John Carey, Alexander Bisdorff, Maurizio Versino, Stefan Evers, and David Newman-Toker. 2012. “Vestibular Migraine: Diagnostic Criteria.” Journal of Vestibular Research: Equilibrium & Orientation 22 (4): 167–72. https://doi.org/10.3233/VES-2012-0453.

Lionel, Anath C., Gregory Costain, Nasim Monfared, Susan Walker, Miriam S. Reuter, S. Mohsen Hosseini, Bhooma Thiruvahindrapuram, et al. 2018. “Improved Diagnostic Yield Compared with Targeted Gene Sequencing Panels Suggests a Role for Whole-Genome Sequencing as a First-Tier Genetic Test.” Genetics in Medicine 20 (4): 435–43. https://doi.org/10.1038/gim.2017.119.

Lopez-Escamez, Jose A., John Carey, Won Ho Chung, Joel A. Goebel, Måns Magnusson, Marco Mandalà, David E. Newman-Toker, et al. 2015. “Diagnostic Criteria for Menière’s Disease.” Journal of Vestibular Research: Equilibrium and Orientation 25 (1): 1–7. https://doi.org/10.3233/VES-150549.

Martín-Sierra, Carmen, Alvaro Gallego-Martinez, Teresa Requena, Lidia Frejo, Angel Batuecas-Caletrío, and Jose A. Lopez-Escamez. 2017. “Variable Expressivity and Genetic Heterogeneity Involving DPT and SEMA3D Genes in Autosomal Dominant Familial Meniere’s Disease.” European Journal of Human Genetics: EJHG 25 (2): 200–207. https://doi.org/10.1038/ejhg.2016.154.

Martín-Sierra, Carmen, Teresa Requena, Lidia Frejo, Steven D. Price, Alvaro Gallego-Martinez, Angel Batuecas-Caletrio, Sofía Santos-Pérez, Andrés Soto-Varela, Anna Lysakowski, and Jose A. Lopez-Escamez. 2016. “A Novel Missense Variant in PRKCB Segregates Low-Frequency Hearing Loss in an Autosomal Dominant Family with Meniere’s Disease.” Human Molecular Genetics 25 (16): 3407–15. https://doi.org/10.1093/hmg/ddw183.

Mašindová, Ivica, Andrea Šoltýsová, Lukáš Varga, Petra Mátyás, Andrej Ficek, Miloslava Hučková, Martina Sůrová, et al. 2015. “MARVELD2 (DFNB49) Mutations in the Hearing Impaired Central European Roma Population - Prevalence, Clinical Impact and the Common Origin.” PLoS ONE 10 (4). https://doi.org/10.1371/journal.pone.0124232.

Meyer, Michael J., Juan Felipe Beltrán, Siqi Liang, Robert Fragoza, Aaron Rumack, Jin Liang, Xiaomu Wei, and Haiyuan Yu. 2018. “Interactome INSIDER: A Structural Interactome Browser for Genomic Studies.” Nature Methods 15 (2): 107–14. https://doi.org/10.1038/nmeth.4540.

Mijovic, Tamara, Anthony Zeitouni, and Inés Colmegna. 2013. “Autoimmune Sensorineural Hearing Loss: The Otology–Rheumatology Interface.” Rheumatology 52 (5): 780–89. https://doi.org/10.1093/rheumatology/ket009.

Mitchell, Kevin J. 2012. “What Is Complex about Complex Disorders?” Genome Biology 13 (1): 237. https://doi.org/10.1186/gb-2012-13-1-237.

Miyasaka, Yuki, Hiroshi Shitara, Sari Suzuki, Sachi Yoshimoto, Yuta Seki, Yasuhiro Ohshiba, Kazuhiro Okumura, et al. 2016. “Heterozygous Mutation of Ush1g/Sans in Mice Causes Early-Onset Progressive Hearing Loss, Which Is Recovered by Reconstituting the Strain-Specific Mutation in Cdh23.” Human Molecular Genetics 25 (10): 2045–59. https://doi.org/10.1093/hmg/ddw078.

Morales Angulo, C., R. Gómez Castellanos, J. García Mantilla, J. T. Bezos Capelastegui, and F. Carrera. 2003. “[Epidemiology of Menière’s disease in Cantabria].” Acta Otorrinolaringologica Espanola 54 (9): 601–5.

Paparella, Michael M., and Matthew S. Griebie. 1984. “Bilaterality of Meniere’s Disease.” Acta Oto-Laryngologica 97 (3–4): 233–37. https://doi.org/10.3109/00016488409130984.

Requena, T., J. M. Espinosa-Sanchez, S. Cabrera, G. Trinidad, A. Soto-Varela, S. Santos-Perez, R. Teggi, et al. 2014. “Familial Clustering and Genetic Heterogeneity in Meniere’s Disease.” Clinical Genetics 85 (3): 245–52. https://doi.org/10.1111/cge.12150.

Requena, Teresa, Sonia Cabrera, Carmen Martín-Sierra, Steven D. Price, Anna Lysakowski, and José A. Lopez-Escamez. 2015. “Identification of Two Novel Mutations in FAM136A and DTNA Genes in Autosomal-Dominant Familial Meniere’s Disease.” Human Molecular Genetics 24 (4): 1119–26. https://doi.org/10.1093/hmg/ddu524.

Requena, Teresa, Juan M. Espinosa-Sanchez, and Jose A. Lopez-Escamez. 2014. “Genetics of Dizziness: Cerebellar and Vestibular Disorders.” Current Opinion in Neurology 27 (1): 98–104. https://doi.org/10.1097/WCO.0000000000000053.

Requena, Teresa, Irene Gazquez, Antonia Moreno, Angel Batuecas, Ismael Aran, Andres Soto-Varela, Sofia Santos-Perez, et al. 2013. “Allelic Variants in TLR10 Gene May Influence Bilateral Affectation and Clinical Course of Meniere’s Disease.” Immunogenetics 65 (5): 345–55. https://doi.org/10.1007/s00251-013-0683-z.

Ruiz-Pesini, Eduardo, Marie T. Lott, Vincent Procaccio, Jason C. Poole, Marty C. Brandon, Dan Mishmar, Christina Yi, James Kreuziger, Pierre Baldi, and Douglas C. Wallace. 2007. “An Enhanced MITOMAP with a Global MtDNA Mutational Phylogeny.” Nucleic Acids Research 35 (Database issue): D823–828. https://doi.org/10.1093/nar/gkl927.

Shearer, A. Eliot, Michael S. Hildebrand, and Richard JH Smith. 1993. “Hereditary Hearing Loss and Deafness Overview.” In GeneReviews(r), edited by Margaret P. Adam, Holly

H. Ardinger, Roberta A. Pagon, Stephanie E. Wallace, Lora JH Bean, Karen Stephens, and Anne Amemiya. Seattle (WA): University of Washington, Seattle. http://www.ncbi.nlm.nih.gov/books/NBK1434/.

Shearer, A. Eliot, and Richard J.H. Smith. 2015. “Massively Parallel Sequencing for Genetic Diagnosis of Hearing Loss: The New Standard of Care.” Otolaryngology--Head and Neck Surgery?: Official Journal of American Academy of Otolaryngology-Head and Neck Surgery 153 (2): 175–82. https://doi.org/10.1177/0194599815591156.

Sloan-Heggen, Christina M., Amanda O. Bierer, A. Eliot Shearer, Diana L. Kolbe, Carla J. Nishimura, Kathy L. Frees, Sean S. Ephraim, et al. 2016. “Comprehensive Genetic Testing in the Clinical Evaluation of 1119 Patients with Hearing Loss.” Human Genetics 135 (4): 441–50. https://doi.org/10.1007/s00439-016-1648-8.

The Molecular Otolaryngology and Renal Research Laboratories, The University of Iowa.n.d. “Deafness Variation Database.” Accessed June 30, 2018. http://deafnessvariationdatabase.org/.

Weber, Megan L., Hong-Yuan Hsin, Ersan Kalay, Dana S. BroŽková, Takehiko Shimizu, Merve Bayram, Kathleen Deeley, et al. 2014. “Role of Estrogen Related Receptor Beta (ESRRB) in DFN35B Hearing Impairment and Dental Decay.” BMC Medical Genetics 15 (July): 81. https://doi.org/10.1186/1471-2350-15-81.

Weil, Dominique, Aziz El-Amraoui, Saber Masmoudi, Mirna Mustapha, Yoshiaki Kikkawa, Sophie Lainé, Sedigheh Delmaghani, et al. 2003. “Usher Syndrome Type I G (USH1G) Is Caused by Mutations in the Gene Encoding SANS, a Protein That Associates with the USH1C Protein, Harmonin.” Human Molecular Genetics 12 (5): 463–71.

Yang, Tao, Hilmar Vidarsson, Sandra Rodrigo-Blomqvist, Sally S. Rosengren, Sven Enerback, and Richard J. H. Smith. 2007. “Transcriptional Control of SLC26A4 Is Involved in Pendred Syndrome and Nonsyndromic Enlargement of Vestibular Aqueduct (DFNB4).” American Journal of Human Genetics 80 (6): 1055–63. https://doi.org/10.1086/518314.

